# Microbial community analyses of composts are influenced by particle size fraction and DNA extraction method

**DOI:** 10.1101/2025.10.15.682543

**Authors:** Anja Logo, Tabea Koch, Monika Maurhofer, Thomas Oberhänsli, Barbara Thürig, Franco Widmer, Pascale Flury, Johanna Mayerhofer

**Author notes:** Corresponding authors: *Email addresses:* (Anja Logo), (Tabea Koch), (Thomas Oberhänsli), (Barbara Thürig), (Pascale Flury), (Johanna Mayerhofer). These authors contributed equally to the work (Pascale Flury, Johanna Mayerhofer).

## Abstract

Composting plays a key role in sustainable agriculture by converting organic waste into a valuable soil conditioner. The process is driven by complex microbial communities, whose characterization is essential for optimizing the composting process and compost quality. Molecular techniques such as amplicon sequencing are commonly used for this purpose. However, sampling procedures and DNA extraction methods, key steps in the sequencing workflow, vary often across studies, challenging comparability.

We investigated two aspects of sampling preparation that may influence compost microbial analyses. For DNA extraction, often fine fractions (*<*2 mm) are used. However, compost has a heterogeneous structure, including coarse particles. To assess the effect of particle size, we separately sequenced bacterial and fungal communities of the fine (0–2 mm) and coarse (2–10 mm) fractions of three composts. In addition, DNA was extracted using a carboxyl affinity-based magnetic method and a silanol affinity-based filter method to evaluate the impact of the extraction technique.

We found that the coarse fraction had higher bacterial richness and a distinct bacterial and fungal community structure compared to the fine fraction. DNA extraction method also influenced bacterial community profiles, with the magnetic bead method improving coverage, particularly for *Bacillota*. Although the effects of particle size and extraction method were small compared to general diversity among composts, we recommend including coarse particles in sequencing analyses and using standardized DNA extraction protocols, especially for studies aiming at high-resolution community analyses.

**Graphical abstract:** 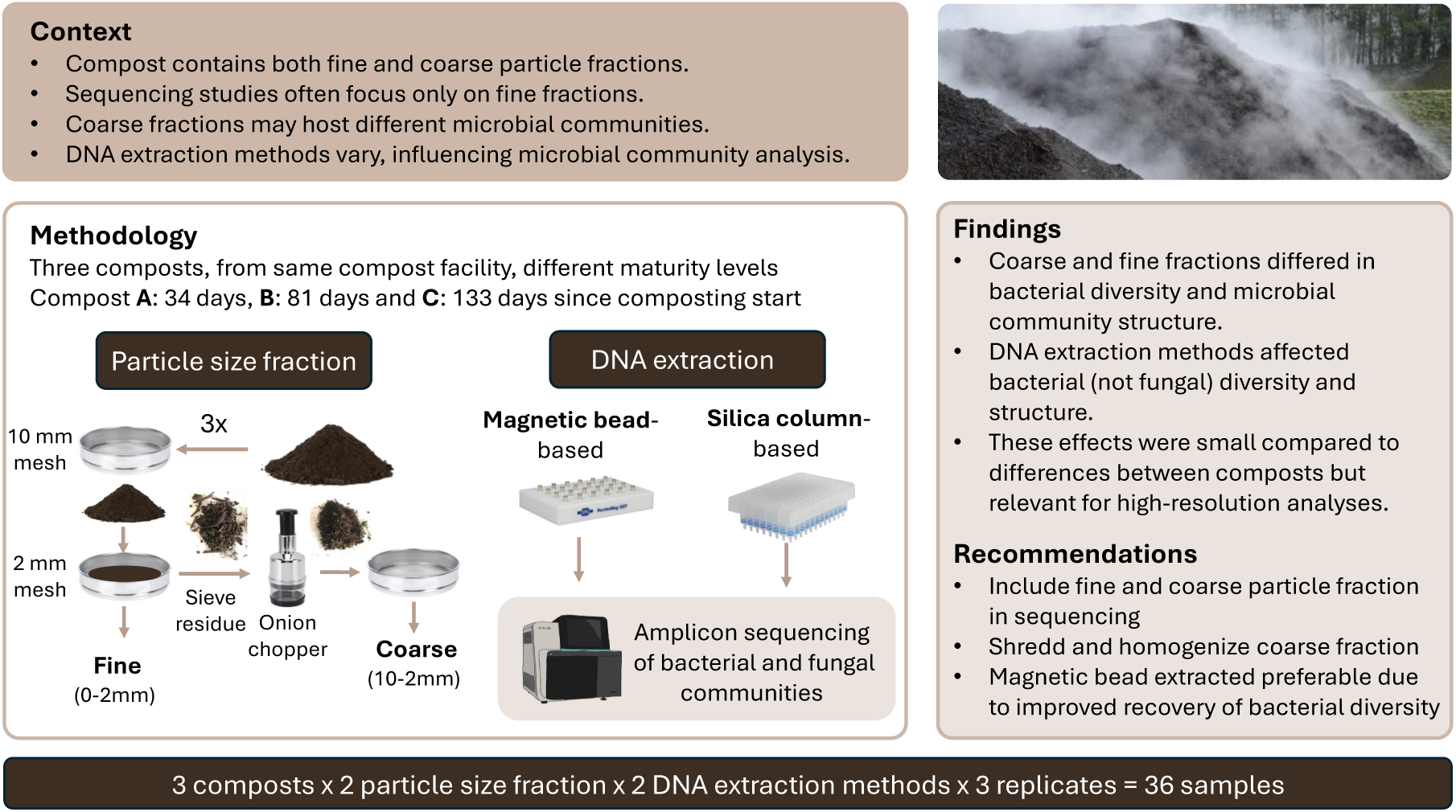

## 1. Introduction

Composting, the aerobically decomposition of organic materials, is not only a sustainable way to recycle organic waste, but the application of compost can also promote soil and plant health (Bonanomi et al., 2010; De Corato, 2020; Aguilar-Paredes et al., 2023). Together with physicochemical parameters the composition of the microbial communities changes along the composting process and plays an important role for the composting process itself, as well as for the quality of the finished compost (Tian et al., 2013). A better understanding of microbial communities and their succession can help to optimize the composting process and enhance compost properties beneficial for plants and soil (Lutz et al., 2020; Luo et al., 2023).

Molecular methods, particularly amplicon sequencing, have been widely used to study compost microbial communities (Parab et al., 2023), i.e., to investigate microbial succession along the composting process (Chang et al., 2021), to define core compost microbiomes (Wang et al., 2020) and to identify bioindicators for compost maturity (EstrellaGonzález et al., 2020), as well as for plant beneficial properties, particularly disease-suppressive activity (Yu et al., 2015; Blaya et al., 2016; Scotti et al., 2020; Mayerhofer et al., 2021; Logo et al., 2025). However, the methodological approaches used to sequence microbial communities in compost vary between individual studies (Parab et al., 2023). This makes it difficult to compare results of studies and conduct meta-studies, which are particularly important given the high complexity of microbial communities in compost and the often low number of observations (e.g. compost types).

Compost has a heterogeneous structure, ranging from fine, highly decomposed particles to large, woody, less decomposed particles. While most studies report the amount of compost used for DNA extraction, particle size is rarely specified. Typically, small sample sizes (200-1,500 mg) are used (Scotti et al., 2020; Mayerhofer et al., 2021) suggesting that predominantly fine particles are sampled. In contrast, for analyses, such as for physicochemical characterization or plant growth assays in pots larger sample sizes are required, which usually contain both coarse and fine particles (Logo et al., 2025).

However, it remains unclear whether the microbial communities of fine and coarse particles are similar and whether microbial communities of composts can be accurately represented by the fine fraction alone. In soils, many microorganisms show preferences for specific particle size fractions due to distinct micro-environments shaped by water availability, nutrient accessibility, and organic matter composition (Sessitsch et al., 2001; Hemkemeyer et al., 2018, 2019). In compost, physicochemical properties also differ between fine and coarse particle fractions (Ĺopez et al., 2002). Fine fractions (*<*2 mm) generally had lower salinity and C:N ratios, but were more mature than coarse fractions (*>*2 mm). Another study showed, that finer fractions contained higher amounts of available nutrients (Hanc and Dreslova, 2016). Moreover, physicochemical properties of composts correlated with microbial community composition (Neher et al., 2013; Wei et al., 2018; Wang et al., 2020). In addition, fungal communities colonized particularly lignin-rich, structurally complex substrates (Ryckeboer et al., 2003), which are more commonly found in coarse particles (De Bertoldi et al., 1983). Building on these findings, we hypothesize that microbial communities in compost differ between fine and coarse particle size fractions.

Another important step in the sample preparation procedure is the DNA isolation from compost, which is challenging due to high levels of humic acids in composts that can co-precipitate with DNA and can inhibit PCR reactions (Matheson et al., 2010). To address this, compost-specific liquid-phase extraction methods have been developed (LaMontagne et al., 2002; Howeler et al., 2003). Nevertheless, compost studies commonly use commercial soil DNA kits, primarily silanol affinity-based filter methods (column-based) (Parab et al., 2023). Recently, carboxyl affinity-based magnetic bead systems (magnetic bead-based) have gained popularity due to their compatibility with automated, high-throughput workflows (Ye and Lei, 2023). Yet, it is unknown whether the choice between column-based and magnetic bead-based extraction methods affects compost microbial community analyses.

In this study, we examined the effects of (I) particle size fraction (fine vs. coarse) and (II) DNA extraction method (column vs. magnetic bead-based) on bacterial and fungal community analyses using amplicon sequencing. We analyzed three composts, all produced by the same composting protocol but differing in maturation time. Our findings contribute to identifying potential methodological biases and provide recommendations for standardizing sampling and DNA extraction approaches in compost microbial community analyses.

## 2. Materials and Methods

### 2.1. Compost sample preparation

Compost samples were collected from three compost piles at the same composting facility in Northwestern Switzerland on July 18, 2022. All three composting processes used the same feedstock composition: 75% green waste from gardeners and the municipality, 20% mature compost, and 5% soil. The composting process was carried out in small triangular piles (1 m height, 1.5 m width), which were turned two to three times per week during the thermophilic phase. The three compost samples (referred to as A, B, and C) differed in the number of days passed since the start of the composting process (i.e., maturity). Sample A was collected after the completion of the thermophilic phase, 34 days after beginning of the composting process. At this composting facility, the compost undergoes further maturation in large piles with regular ventilation. Sample B was taken from one of these maturation piles (81 days after composting start). Afterwards, composts are sieved through a 15 mm-mesh and stored in a covered area. Sample C was taken from this storage area 133 days after the start of the composting process. The sampling procedure involved taking subsamples at a depth of 20 to 30 cm from three to five different locations in a pile and pooling them. The three composts A, B and C were also part of a larger study in which they are named 11, 12 and 13 (Logo et al., 2025).

Of each of the collected composts two fractions were generated to study differences in microbial communities with regard to particle size (Figure 1). A handful of compost was sieved first through a 10 mm-mesh and then through a 2 mm-mesh, which defined the fine particle size fraction (0-2 mm). The material retained on top of the 2 mm sieve was chopped using an onion chopper and sieved again through the 2 mm-mesh. This fraction was considered the coarse particle fraction (2-10 mm). For each of the three composts three replicate samples were prepared per size fraction. Of these samples, two portions of 200 mg were transferred each to a 2 mL screw cap tube containing 1.5 g ceramic beads (SiLibeads Type ZY S 0.4-0.6 mm, Sigmund Lindner, Germany). To one of the two samples, 1 mL NucleoMag lysis buffer was added. Samples were stored at-20°C until DNA extraction.

**Figure 1:**
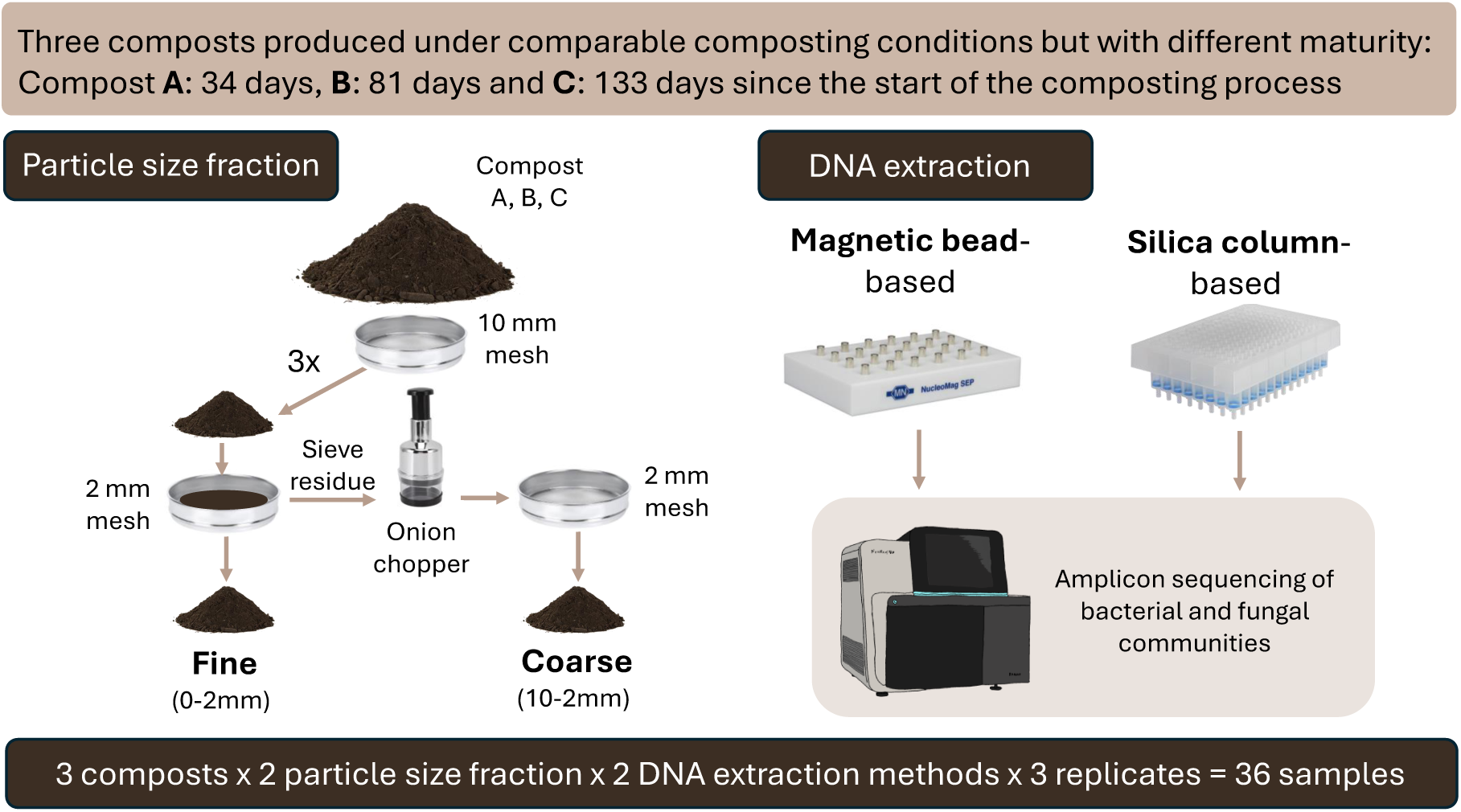
Experimental set up

### 2.2. DNA extraction

The samples with the NucleoMag lysis buffer were extracted using a carboxyl affinity-based magnetic bead DNA extraction kit (NucleoMag DNA Microbiome for DNA purification from soil, stool, and biofilm), hereafter referred to as magnetic bead-based DNA extraction. The second sample was extracted using a silanol affinity-based filter DNA extraction kit (NucleoSpin 96 Soil Extraction Kit), hereafter referred to as column-based DNA extraction. Both kits were produced by Macherey-Nagel (Düren, Germany). DNA extraction was performed according to the manufacturer’s instructions with a few adaptations. Samples were homogenized using a TissueLyser (Qiagen, The Netherlands) at 30 Hz for 4 min. For the bead-based extraction, DNA on magnetic beads was dried at 70°C for 30 min to remove residual ethanol, then eluted in 100 *µ*L of 0.1× TE buffer (1 mM Tris, 0.1 mM Na_2_EDTA, pH 8.0). For the column-based extraction, 1 mL of lysis buffer without SX Enhance was used for cell lysis, and DNA was eluted in 200 *µ*L in two steps. DNA was quantified using Quant-iT PicoGreen dsDNA Assay kit (Invitrogen, Waltham, MA, USA) with a Cary Eclipse fluorescence spectrophotometer (Varian, Palo Alto, CA, USA).

### 2.3. Quantitative real-time PCR

The same marker genes were used for quantitative real-time PCR (qPCR) and for the PCR for amplicon sequencing (see Section 2.4): for bacteria, the variable region 3 and 4 of the 16S rRNA marker gene (16S); and for fungi the internal transcribed spacer regions (ITS2). However, for qPCR, primers with fewer degenerate bases were selected to increase amplification efficiency. All primer pairs are listed in Table S1.

For qPCR, DNA extracts were diluted 1:20 for bacteria and 1:10 for fungi using deionized, autoclaved H_2_O. The qPCR master mix consisted of 1x qPCRBIO SyGreen Blue Mix (PCR Biosystems, London, UK), 1.8 *µ*M of each primer, and 0.3 mg L^-1^ bovine serum albumin (BSA). The total reaction volume was 10 *µ*L, including 2 *µ*L of the diluted DNA extract. The qPCR assay was run in triplicates using a CFX Opus 384 Real-Time PCR system (BioRad, Hercules, CA, USA). Thermal cycling conditions started with enzyme activation at 95°C for 3 min, followed by 40 cycles of degeneration at 95°C for 10 s, annealing at 53°C for 40 s and elongation at 72°C for 1 min. The process was finished with a melt curve from 65 to 95°C increasing by 0.5°C per cycle to verify the specificity of the amplification product. Bacterial and fungal qPCRs were run on separate plates. Amplification curves were analyzed using BioRad CFX Maestro 2.3 and fluorescent threshold was set manually to 600 RFU.

A synthetic double-stranded DNA fragment (gblock, HiFi gene fragments, IDT, Coralville, IA, USA) containing a single copy of the target gene sequence was used to prepare the standard dilution series ranging from 10^7^ to 10^4^ or 10^3^ copies per *µ*L for bacteria and fungi, respectively. Standard curves, which are shown in Figure S1, were used to calculate bacterial and fungal copy numbers from Ct values. Amplification efficiency was at 79.1% (R^2^ = 0.996) for bacteria and 95% (R^2^ = 0.997) for fungi. For all samples bacterial and fungal copy numbers were normalized to 1 ng of DNA (DNA concentration and non-normalized copy numbers in Table S2).

### 2.4. Amplicon sequencing

For PCR amplification, DNA extracts were diluted to 5 ng *µ*L^-1^. Each PCR reaction contained 20 ng DNA, 1 *µ*M of each primer, 0.2 mM dNTPs (Promega, Madison, IA, USA), 2.5 mM MgCl_2_, 0.6 mg mL^-1^BSA, GoTaq Hotstart Flexi buffer (Promega, Madison, IA, USA), 1.25 units of GoTaq Hotstart G2 polymerase and deionized, autoclaved H_2_O to a final volume of 25 *µ*L. PCR reactions were performed in 96-well plates.

Thermal cycling conditions started with an initial denaturation step at 95°C for 2 min, followed by 30 cycles for bacteria and 33 cycles for fungi, consisting of denaturation at 94°C for 40 s, annealing at 56°C for 40 s and elongation at 72°C for 1 min. PCR reactions were run in quadruplets and pooled after quality control on an Agarose gel. PCR product concentrations were quantified as described for genomic DNA and diluted to 20 ng *µ*L^-1^. Library preparation included tagging with TruSeq indices, normalization of read counts from an Illumina MiniSeq run (Illumina Inc., San Diego, CA, USA), and sequencing on the Illumina NextSeq2000 platform. Paired-end sequencing was performed with 300 bp long reads at the Functional Genomics Center Zurich (University of Zurich / ETH Zurich, Switzerland)).

### 2.5. Bioinformatics and quality control

Quality control, ASV inference and taxonomic sequence classification was performed using a pipeline adapted from Frey et al. (2016) and Mayerhofer et al. (2021). The pipeline, particularly the commands and package versions used, is detailed in our previous publication (Logo et al., 2025). Briefly, primers were trimmed using *cutadapt*, poly-G tails were trimmed and sequences with low complexity (cut-off 60) and shorter than 150 bps were removed using *fastp* version 0.23.4 (Chen et al., 2018). Then forward and reverse sequences were merged, quality filtered and dereplicated in *vsearch*. Sequences were clustered to amplicon sequence variants (ASVs) using *UNOISE*. Chimera were removed with *UCHIME*. Ribosomonal signature was searched using *Metaxa2* for bacteria and *ITSx* for fungi and sequences without discarded. ASVs were taxonomically classified using the SILVA Database for bacteria (Release SSU 138.2; Quast et al. (2012)) and UNITE database for fungi (version 9.0; Abarenkov et al. (2024)). Non-bacterial sequences and non-fungal sequences were removed. The number of sequences and ASVs at different filtering steps are summarized in Table S3.

### 2.6. Statistical analyses

All analyses and visualizations were performed in R (v.4.3.1, R Core Team (2023)) using Rstudio (v.2023.06) and the *tidyverse* packages (Wickham et al., 2019). Statistical significance was defined at *p <*0.05. Microbial community analyses were performed using the vegan package (Oksanen et al., 2022). Due to variable sequencing depth (Figure S2), samples were rarefied to the lowest observed depth (96,235 sequences for bacteria; 88,625 for fungi), repeated 100 times using”*rarefy*”. Richness and evenness were calculated using”*specnumber*” and”*diversity*” and Bray–Curtis dissimilarities using”*vegdist*” for all 100 subsampled ASV tables and averaged.

To account for the hierarchical sampling design (Figure 1), linear mixed models (”*lmer*”, *lmerTest*) were used to test for differences in alpha diversity and microbial quantity. Compost, particle size, and extraction method (including interactions) were treated as fixed effects, and”compost-replicate” as a random effect. If random variance was negligible linear models (”*lm*”) were used. Model assumptions were checked via residual plots. Type III ANOVA tables were generated using”*Anova*” (*car*) for mixed models and”*anova*” for linear models. Estimated marginal means (EMMs) were calculated using *emmeans* (Lenth, 2024), with S^̌^idák correction for pairwise comparisons.

Community structure difference (Bray–Curtis) were tested via PERMANOVA (”*adonis2*”, *vegan*) with permutations constrained within”compost-replicates”. Community compositions were visualized through Principal Coordinate Analysis (PCoA) using”*cmdscale*”.

To identify treatment-associated ASVs (coarse vs. fine; beads vs. column), we conducted two analyses. First, we identified treatment-specific ASVs, defined as those exclusively present in one treatment level. Only robustly detected ASVs were considered (present in at least two of three technical replicates in at least one sample). Second, we identified ASVs consistently enriched across particle sizes or extraction methods, based on absolute abundances. These were calculated by multiplying relative abundances (from unrarefied read counts) by the measured bacterial or fungal quantity. For each ASV, we calculated log_2_ fold change and mean relative abundance in the respective compost. ASVs with relative abundance below 0.01% in all replicates or log_2_ fold change below 1 were excluded. Treatment-associated ASVs were identify separately per compost and then compared.

## 3. Results

We assessed how microbial community analyses of composts were influenced by (I) extracting DNA from fine (0-2 mm) vs. coarse (2-10 mm) particle fractions and (II) using either a column-based or magnetic bead-based DNA extraction method. The data set consisted of three composts (named A-C), each represented by three technical replicates, all collected on the same day and from the same producer, but with different degrees of maturity (C *>*B *>*A). Overall, sequencing of the 36 samples yielded 5,052,447 highquality sequences of bacteria and 4,268,339 of fungi (an average of 140,346 and 118,565 and a SD of 32,161 and of 13,396 per sample, respectively). The flattening of rarefaction curves and Good’s coverage exceeding 99% in all samples indicate that sequencing depth was sufficient (Figure S3).

The sequences clustered into 4,871 bacterial and 851 fungal Amplicon Sequences variants (ASVs). Of these, 75.4% of bacterial and 45.8% fungal ASVs were found in all three composts, representing 98.8% of all bacterial and 99.6% of fungal sequences. Thus, the most common bacteria and fungi were found in all composts, but differed in their abundance between composts (Figure S4).

### 3.1. Bacterial diversity estimates were increased when DNA was extracted from the coarse particle fraction and with magnetic beads

Quantitative PCR analysis showed that the copy numbers of bacterial and fungal ribosomal marker genes per ng of total DNA varied with compost, particle size fraction and DNA extraction method (Figure 2A and B; statistical test values in Table 1). Mean bacterial gene copy number increased by 66.7% from compost A to compost C, reflecting increasing compost maturity, while mean fungal copy number decreased slightly by 4.2%. Additionally, both bacterial and fungal quantities were higher in the fine particle fraction, with 35.1% and 43.8% higher mean values, respectively. Similarly, higher quantities were observed in samples extracted using magnetic beads, with 33.0% higher mean values for bacteria and 14.1% for fungi.

**Figure 2:**
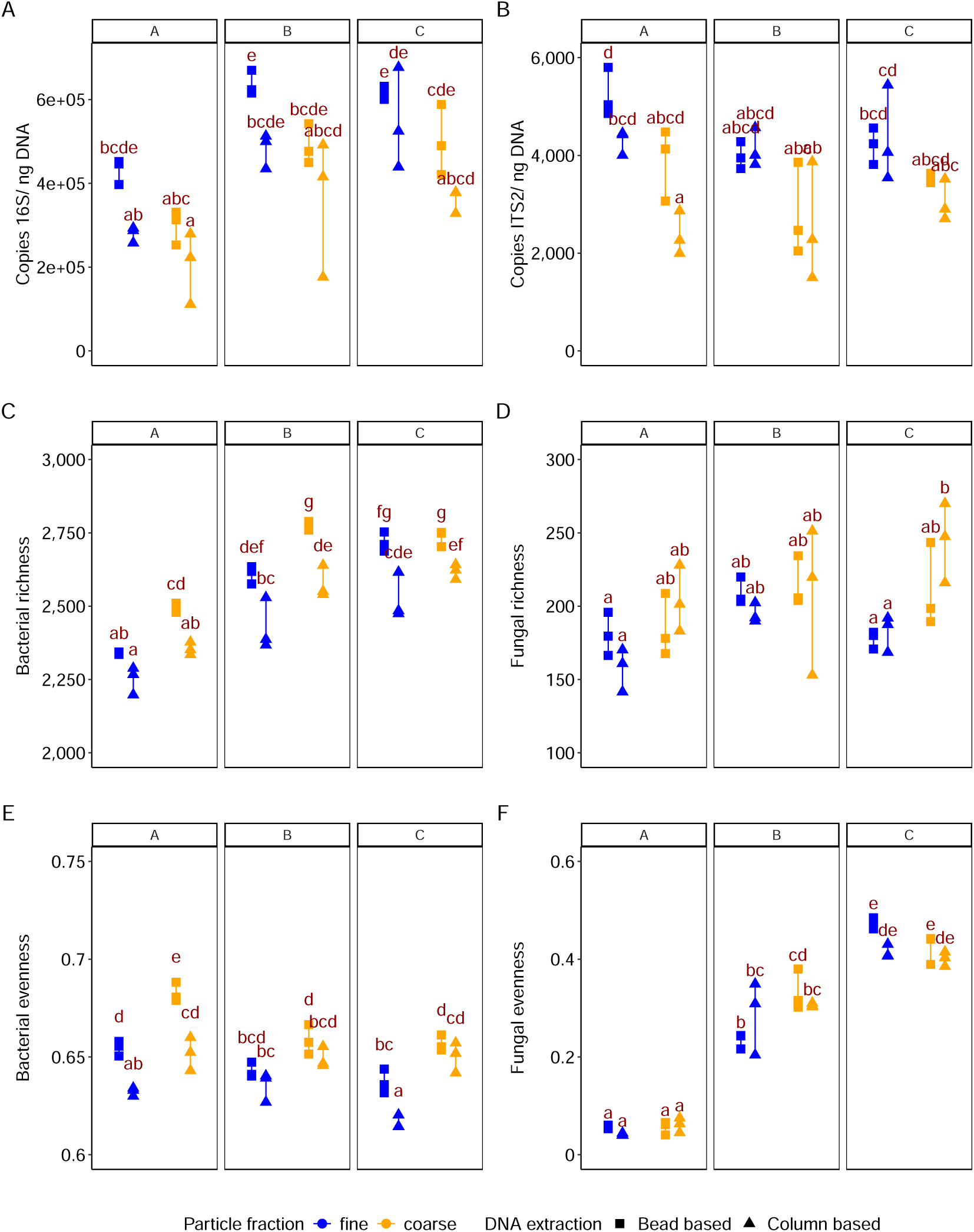
F**u**ngal **quantity and bacterial quantity and diversity differed between particle size fractions and DNA extraction methods.** Effects of compost type, particle size fraction, DNA extraction method, and their interactions on bacterial and fungal (A, B) absolute quantity, i.e., ribosomal gene copy numbers (C, D) ASV richness, and (E, F) Shannon evenness of ASVs were tested using mixed linear models with “compost-replicate” as a random factor, or standard linear models when the random effect was negligible. F-values and p-values for fixed factors are shown in Table 1 and full Type III ANOVA results can be found in the Supplementary Materials (Table S6 and S7). Treatments sharing the same red letters within a subplot (A-F) do not differ significantly (post-hoc S^̌^id‘ak corrected pairwise comparisons).

**Table 1:**
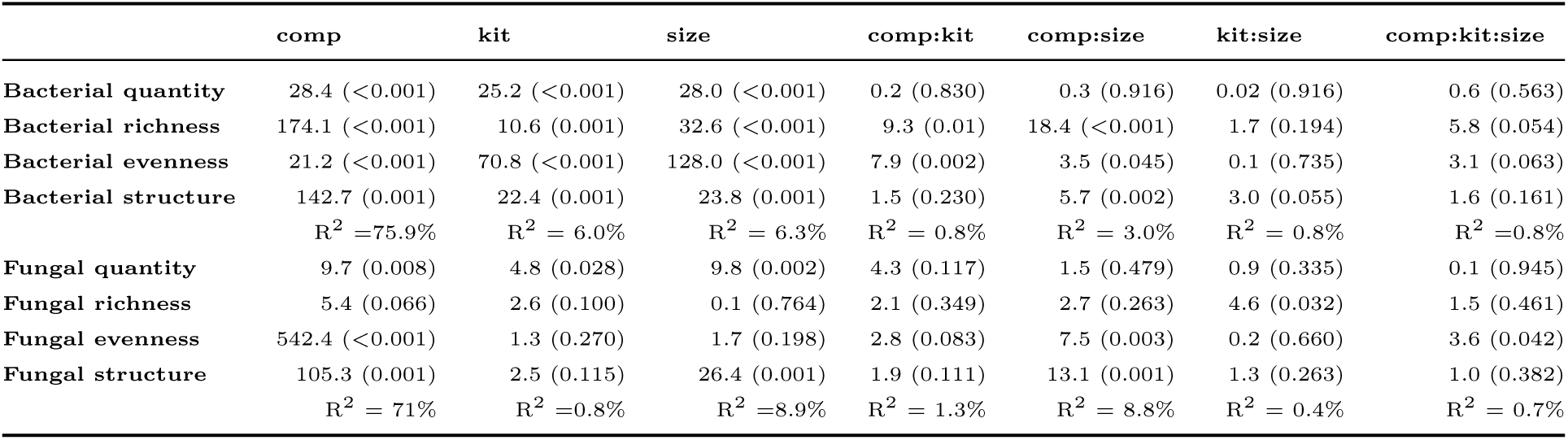
Summary of statistical analysis of the effects of compost (comp), particle size (size) and DNA extraction (kit) on bacterial and fungal communities. For bacterial quantity, bacterial evenness and fungal evenness a standard linear model was applied. For bacterial richness, fungal quantity and fungal richness a mixed-linear model with the compost-replicate as a random factor was applied. Difference in community structure were tested with a PERMANOVA with permutations restricted by compost-replicate. Shown are the corresponding F-values / Chi-square and Pseudo-F values and in brackets the p-values. For the community structures, also R^2^ are shown. Full model outputs are provided in the Tables S6, S7 and S8.

|  | comp | kit | size | comp:kit | comp:size | kit:size | comp:kit:size |
| --- | --- | --- | --- | --- | --- | --- | --- |
| **Bacterial quantity** | 28.4 (<0.001) | 25.2 (<0.001) | 28.0 (<0.001) | 0.2 (0.830) | 0.3 (0.916) | 0.02 (0.916) | 0.6 (0.563) |
| **Bacterial richness** | 174.1 (<0.001) | 10.6 (0.001) | 32.6 (<0.001) | 9.3 (0.01) | 18.4 (<0.001) | 1.7 (0.194) | 5.8 (0.054) |
| **Bacterial evenness** | 21.2 (<0.001) | 70.8 (<0.001) | 128.0 (<0.001) | 7.9 (0.002) | 3.5 (0.045) | 0.1 (0.735) | 3.1 (0.063) |
| **Bacterial structure** | 142.7 (0.001)<br>$R^2 = 75.9\%$ | 22.4 (0.001)<br>$R^2 = 6.0\%$ | 23.8 (0.001)<br>$R^2 = 6.3\%$ | 1.5 (0.230)<br>$R^2 = 0.8\%$ | 5.7 (0.002)<br>$R^2 = 3.0\%$ | 3.0 (0.055)<br>$R^2 = 0.8\%$ | 1.6 (0.161)<br>$R^2 = 0.8\%$ |
| **Fungal quantity** | 9.7 (0.008) | 4.8 (0.028) | 9.8 (0.002) | 4.3 (0.117) | 1.5 (0.479) | 0.9 (0.335) | 0.1 (0.945) |
| **Fungal richness** | 5.4 (0.066) | 2.6 (0.100) | 0.1 (0.764) | 2.1 (0.349) | 2.7 (0.263) | 4.6 (0.032) | 1.5 (0.461) |
| **Fungal evenness** | 542.4 (<0.001) | 1.3 (0.270) | 1.7 (0.198) | 2.8 (0.083) | 7.5 (0.003) | 0.2 (0.660) | 3.6 (0.042) |
| **Fungal structure** | 105.3 (0.001)<br>$R^2 = 71\%$ | 2.5 (0.115)<br>$R^2 = 0.8\%$ | 26.4 (0.001)<br>$R^2 = 8.9\%$ | 1.9 (0.111)<br>$R^2 = 1.3\%$ | 13.1 (0.001)<br>$R^2 = 8.8\%$ | 1.3 (0.263)<br>$R^2 = 0.4\%$ | 1.0 (0.382)<br>$R^2 = 0.7\%$ |

Alpha diversity differed among composts for both bacteria and fungi, and for bacteria also varied between particle size fractions and DNA extraction methods (Figure 2C-F; statistical test values in Table 1). Mean bacterial richness and mean fungal richness increased by 12.3% and 12.1%, respectively, from compost A to compost C. While mean fungal evenness increased eightfold, mean bacterial evenness decreased by 2.4%. In all three composts, bacterial richness and evenness were higher when DNA was extracted from the coarse particle fraction (mean values higher by 4.6% and 3.4%, respectively) and when using magnetic bead-based extraction method (mean values higher by 6.2% and 2.5%, respectively). However, effect sizes varied among composts, as shown by significant compost × size and compost × kit interactions (Table 1). For fungi, a non-significant trend suggested a higher fungal richness in the coarse fraction, but only when using the column-based DNA extraction method.

In summary, bacterial and fungal quantities, as well as bacterial diversity, varied with compost maturity, particle size fraction, and DNA extraction method, whereas fungal diversity only with compost maturity. DNA extracted from coarse particles yielded lower bacterial and fungal quantities but higher bacterial diversity compared to fine particles. Similarly, samples extracted using magnetic beads showed higher bacterial and fungal quantities, along with higher bacterial diversity, than those extracted with the columnbased method.

### 3.2. Bacterial community structure differed by particle size fraction and DNA extraction method, while fungal structure varied only with particle size

The bacterial and fungal community structure differed strongest among the composts (R^2^_Bacteria_ = 75.9 %, R^2^_Fungi_ = 71%), followed by the particle size fractions (R^2^_Bacteria_ = 6.3 %, R^2^_Fungi_ = 8.9%) and, only for bacteria, among the DNA extraction methods (R^2^ = 6.0%, Figure 3, PERMANOVA test values in Table 1). For bacteria, the effect of the particle size was comparable across composts, with the fine fraction separating from the coarse fraction along the first principle coordinate, the same direction in which also the compost were separated based on maturity. For fungi the differences between particle size fractions increased with compost maturity (C *>*B *>*A). Additionally, while the three technical replicates for bacteria clustered closely together, indicating consistent results, the fungi showed more variability between replicates. This suggests a more heterogeneous distribution of fungal communities.

**Figure 3:**
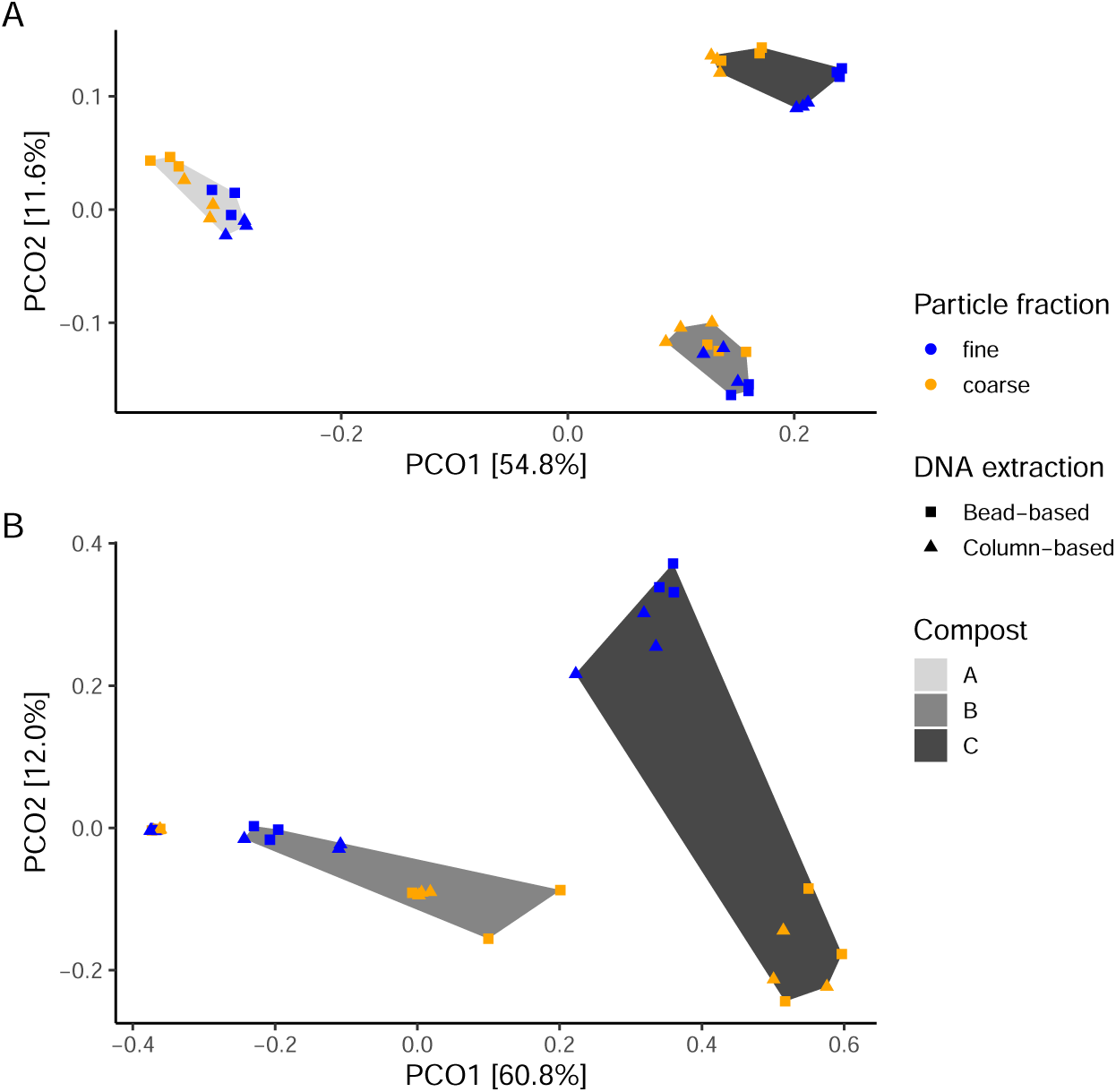
T**h**e **particle size fraction influenced the structure of bacterial and fungal communities, while the DNA extraction method only influenced the bacteria.** The Principle Coordinate Analysis (PCoA) is based on a Bray-Curtis dissimilarity matrix for (A) bacteria and (B) fungi. Treatment effects were tested with a PERMANOVA for which permutations were constrained within”compost-replicates”. Pseudo-F and p-values are indicated in Table 1 and full model outputs in the Table S8.

### 3.3. Particle size and DNA extraction differences resulted from abundance shifts, rather than the presence of specific ASVs

Finally, we aimed to identify specific taxa that differ between particle size fractions or DNA extraction methods. First, we identified ASVs that occurred exclusively in a single treatment level (e.g., present only in the coarse fraction), hereafter referred to as treatment-specific ASVs. Second, we identified ASVs that were consistently enriched in a treatment-level based on their absolute abundances, referred to as treatment-enriched ASVs. Absolute abundances were calculated by multiplying each ASV’s relative abundance (based on unrarefied sequence counts) by the corresponding marker gene copy number estimated by qPCR. Treatment-specific and treatment-enriched ASVs were analyzed separately for each compost and subsequently compared across composts.

Treatment-specific bacterial ASVs ranged from 177 ASVs in column-based extractions of compost B to 323 ASVs in the coarse fraction of compost A (Table S4). In fungi, the number of treatment-specific ASVs ranged from 26 ASVs in the fine fraction of compost A to 77 ASVs in the coarse fraction of compost C (Table S4). However, treatment-specific ASVs were generally low in abundance for both bacteria and fungi, with summed relative abundances across all treatment levels below 0.15%, and were mostly specific to a single compost. The greatest overlap of treatment-specific ASVs occurred in the coarse particle fraction, where eight bacterial and four fungal ASVs were shared among all three composts (Table S4). These results indicate that only few and low abundant ASVs will be entirely missed due the method choice and that treatment-specific ASVs alone are insufficient to explain the observed shifts in microbial community structure associated with particle size fractions and DNA extraction methods.

For the particle size fraction, the highest number of treatment-enriched ASVs was observed in compost C, with 93 bacterial and 5 fungal ASVs enriched in the fine fraction (Table S5). These ASVs accounted for 8.6% and 41.7% of the bacterial and fungal sequences in this compost, respectively. For the DNA extraction method, the highest number of treatment-enriched ASVs was found for compost A, with 103 bacterial ASVs being enriched when DNA was extracted with magnetic beads, accounting together in average for 4.9% of sequences in this compost. In contrast, none of the fungal ASVs differed based on DNA extraction method. Similar to the treatment-specific ASVs, treatment-enriched ASVs were also largely unique to individual composts. Only three bacterial ASVs were consistently enriched across all three composts, however, by different treatments: ASV407 (family *Micromonosporaceae*, in coarse fraction), ASV255 (genus *Opitutus*, in fine fraction), and ASV203 (family *Longimicrobiaceae*, extracted with magnetic beads). For fungi, all treatment-enriched ASVs were compost-specific, with the exception of a single ASV—ASV1 (*Mycothermus thermophilus*)—which was enriched in the fine fraction of both composts B and C.

Thus, the effects of particle size fraction and DNA extraction method on treatment-enriched ASVs, which represented the highest taxonomic resolution, appeared to vary among composts. However, when inspecting the classification of the ASVs to a phylum a pattern emerged: in compost A and C, 58.3% (60 ASVs) and 62.3% (38 ASVs), respectively, of ASVs enriched in samples extracted with magnetic beads were classified within the phylum *Bacillota* (Figure 4). While *Bacillota* was also the most common phylum classification among all ASVs in these two composts (33.7% and 33.6% of the ASVs, respectively), it was still markedly overrepresented among the ASVs enriched in magnetic bead-extracted samples. For particle size fraction, the classification at phylum level of the enriched ASVs did not follow clear taxonomic patterns (Figure 4). Assessing the enriched fungal ASVs, showed that some of the most abundant fungal ASVs differed between particle size fractions (Figure S4). In addition to ASV1 (*Mycothermus thermophilus*), which was the most abundant ASV across all composts and enriched in the fine fraction of composts B and C, ASV2 (family *Chaetomiaceae*) was enriched in the fine fraction of compost B; ASV3 (family *Chaetomiaceae*), ASV7 (genus *Myriococcum*), and ASV9 (family *Chaetomiaceae*) were enriched in the fine fraction of compost C; and ASV6 (genus *Iodophanus*) was enriched in the coarse fraction of compost C.

**Figure 4:**
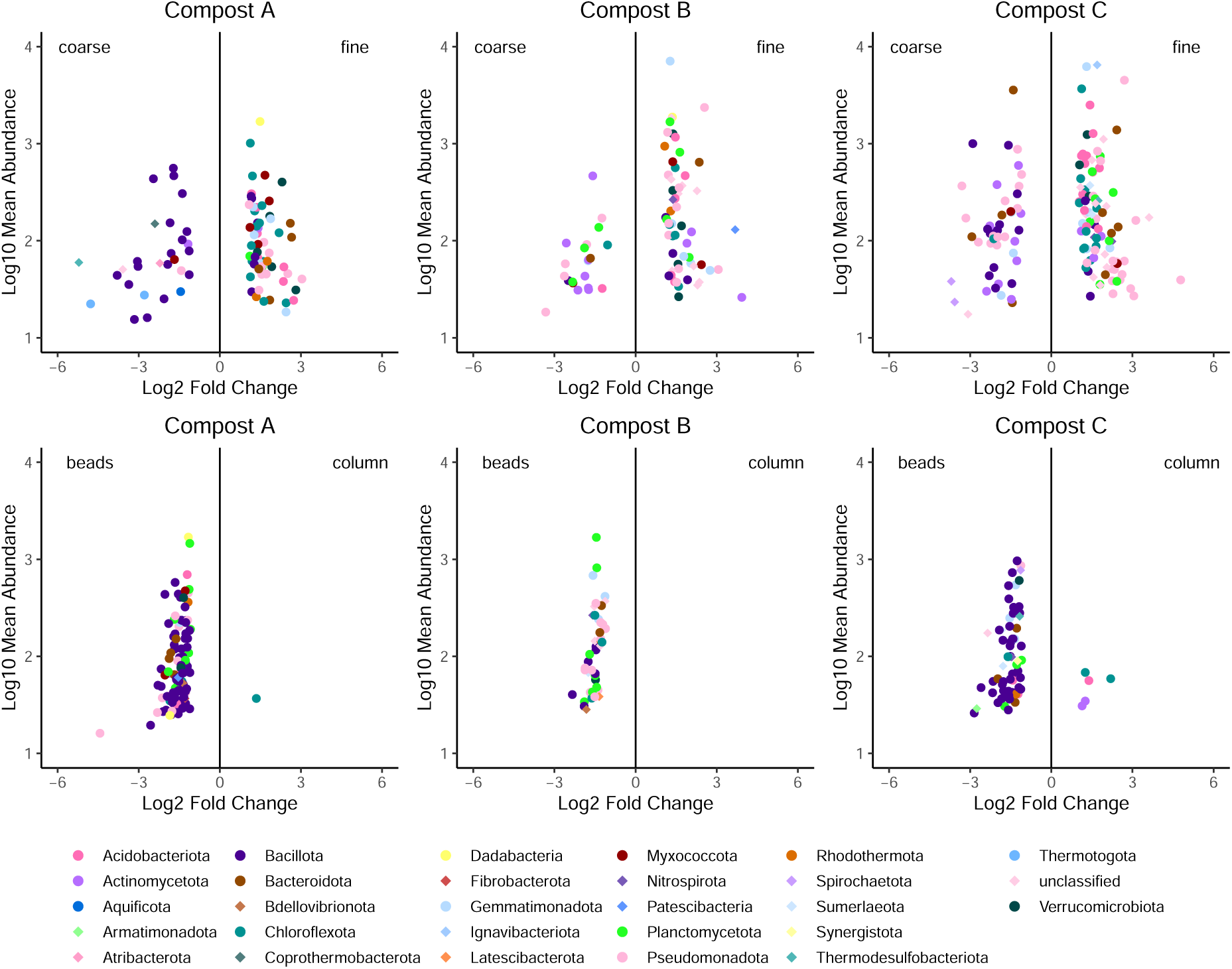
Phylogenetic diverse group of bacteria differed between particle size fractions. Mainly bacteria belonging to the phylum *Bacillota* differed between DNA extraction methods For each ASV, differences in absolute abundance were calculated between samples pairs that differed only in the treatment of interest (particle size or DNA extraction method). An ASV was considered enriched if it was consistently more abundant in one treatment level across all three technical replicates and for both treatment levels of the other factor, i.e. in all six sample pairs. The top plots display results for particle size, while the bottom plots show results for DNA extraction method. Each point represents an ASV, colored by its classification at phylum level. The y-axis shows the mean abundance of the ASV across all samples in the respective compost, and the x-axis shows the mean log_2_ fold change between sample pairs. An ASV is included in the plot when its relative abundance was above 0.01% in at least one replicate and the mean log_2_ fold change exceeded 1.

## 4. Discussion

Differences between composts, reflecting increasing maturity, were the strongest drivers of bacterial and fungal diversity measures and community structure. However, although more subtle, diversity measures and community structure also varied between particle size fractions and, only for bacteria, between DNA extraction methods.

Bacterial richness and fungal evenness increased with compost maturity, consistent with previous findings (Estrella-Gonźalez et al., 2020; Hernández-Lara et al., 2022; Logo et al., 2025). All three compost harbored rich bacterial and fungal communities. However, the least mature compost, compost A, exhibited very low fungal evenness and was dominated by a single ASV, the fungus *Mycothermus thermophilus*. Thus, colonization by diverse fungal communities, which mainly occurs during the cooling and maturation stages (Luo et al., 2023), had only begun for this compost. Differences in community structure between size fractions were observed for bacteria across all composts, although these differences were generally small. In contrast, differences in fungal community structure were limited to composts B and C, where they were notably pronounced. This suggests that particle size fraction, particularly for fungal communities in mature compost, plays a significant role.

Particle size effects were consistent across composts: coarse fractions generally showed higher bacterial diversity (Figure 2), and shifts in community structure followed similar trends for bacteria and fungi (except for compost A; Figure 3). However, taxon-level responses varied, with differences mainly driven by changes in the abundance of taxa, rather than the presence of fraction-specific taxa. This contrasts with soil systems, where specific microbial groups were associated with particular particle sizes (Sessitsch et al., 2001; Hemkemeyer et al., 2018), suggesting that in compost, particle size associations are more dynamic than in soils and may shift with maturity.

Fine compost particles contain more available macro nutrients (Hanc and Dreslova, 2016), potentially promoting fast-growing bacteria, which could explain the higher bacterial quantity. Also, fine particles have physicochemical properties typically associated with increased compost maturity (López et al., 2002). Similarly, bacterial community structure varied with particle size fractions, with differences primarily observed along the first principal coordinate. This coordinate also corresponds to the axis along which compost separates based on maturity (Figure 3). Thus, fine particles tend to harbor more mature bacterial communities, while coarse particles contain less mature ones. In contrast, fungal communities showed the opposite trend for community structure. Moreover, thermophilic fungi, which typically decline with maturity (Wang et al., 2018), were more abundant in the fine fraction. Meanwhile, the coarse fraction of compost C was enriched in the fungus *Iodophanus*, a saprophyte (Richardson, 2001), which typically increase with compost maturity (Xie et al., 2021; Qiao et al., 2025). These results indicate that while both bacteria and fungi respond to particle size, they follow distinct successional patterns.

Pot et al. (2021) compared microbial communities in untreated compost (0–10 mm) and compost with the fine fraction removed (2–10 mm). They found differences in taxa such as *Pirellula*, *Pir5-lineage*, and *Rhodanobacter*, but, relative to differences between compost, the effect of particle size fraction was small. However, their study included composts from five producers, introducing greater variability than our dataset of three composts from a single facility. This similarity is also reflected in the clustering of composts 11, 12, and 13 (equivalent to A–C) in an ordination plot including 37 compost in our previous publication (Figure 4; (Logo et al., 2025)). Therefore, while particle size can influence microbial communities, its effect is relatively minor compared to the variability introduced by different feedstock and facilities. Still, when targeting specific ASVs, e.g. identifying bioindicators, sequencing should include coarse particles. Additionally, since particle size distribution varies with feedstock type (Hanc and Dreslova, 2016), standardizing size through sieving and shredding could enhance comparability across composts.

DNA extraction method also influenced bacterial communities. Magnetic bead extracted samples yielded a higher bacterial richness, driven by increased numbers of ASVs classified to *Bacillota*. Many members of this phylum are Gram-positive, a group generally known to be sensitive to DNA extraction methods in soil and fecal samples (Iturbe-Espinoza et al., 2021; Galla et al., 2024). At the same time, also bacterial quantity was increased in samples extracted with magnetic beads which could partly explain the observed diversity increase, as all DNA extracts were normalized based on the total DNA concentration for sequencing. Higher target gene concentrations can reduce stochastic effects during PCR amplification, potentially resulting in increased diversity estimates (Multinu et al., 2018; Castle et al., 2018). However, it is also possible that DNA purity was higher in the magnetic bead extracted samples, i.e. less humic substances, which can increase PCR efficiency and thus the observed bacterial and fungal quantity. A study comparing DNA extraction methods, including those for soil samples, found that template purity and quantity were key factors affecting diversity estimates (Galla et al., 2024).

For future studies on compost microbial communities, we recommend including both coarse and fine particle fractions, with coarse material shredded and homogenized where necessary. Among the two tested DNA extraction methods, magnetic bead-based extraction is preferred due to its improved recovery of bacterial diversity, possibly through better removal of inhibitors and greater microbial DNA yield. Ultimately, given compost’s heterogeneity, careful and representative sampling remains essential for accurate microbial community analyses.

## CRediT authorship contribution statement

**Anja Logo:** Writing — original draft, Writing — review & editing, Methodology, Visualization, Formal analysis, Conceptualization, Investigation. **Tabea Koch:** Methodology, Investigation. **Monika Maurhofer**: Writing — review & editing, Supervision. **Thomas Oberhänsli**: Project administration, Methodology, Funding acquisition, Writing — review & editing. **Barbara Thürig**: Writing — review & editing, Funding acquisition. **Franco Widmer**: Project administration, Funding acquisition, Conceptualization. **Pascale Flury**: Writing — original draft, Writing — review & editing, Methodology, Conceptualization, Supervision, Funding acquisition. **Johanna Mayerhofer**: Writing — original draft, Writing — review & editing, Formal analysis, Conceptualization, Supervision, Methodology, Funding acquisition.

## Declaration of generative AI and AI-assisted technologies in the writing process

During the preparation of this work the authors used ChatGPT (Version 5, OpenAI) and DeepL in order to fix spelling, grammar and enhancing clarity of the text. After using this tool, the authors reviewed and edited the content as needed and take full responsibility for the content of the published article.

## Acknowledgments

We thank Sonja Reinhard (FiBL) for the support in the lab. We are grateful to the composting facility in North-western Switzerland that provided us the tested composts. We gratefully acknowledge the Functional Genomics Center Zurich (FGCZ) of University of Zurich and ETH Zurich, and in particular Maria Domenica Moccia, for the support in Amplicon sequencing.

## Data availability

The raw sequences have been deposited at the NCBI Sequence Read Archive under the BioProject asccession numbers PRJNA1200637 (bacteria) and PRJNA1201072 (fungi).

