## Supplementary Materials for "Microbial community analyses of composts are influenced by particle size fraction and DNA extraction method"

Additional Material and Methods, Figures and Tables for the paper "*Microbial community analyses of composts are influenced by particle size fraction and DNA extraction method*".

*Authors*: Anja Logo, Tabea Koch, Monika Maurhofer, Thomas Oberhänsli, Barbara Thürig, Franco Widmer, Johanna Mayerhofer & Pascale Flury

*Date*: October 15, 2025

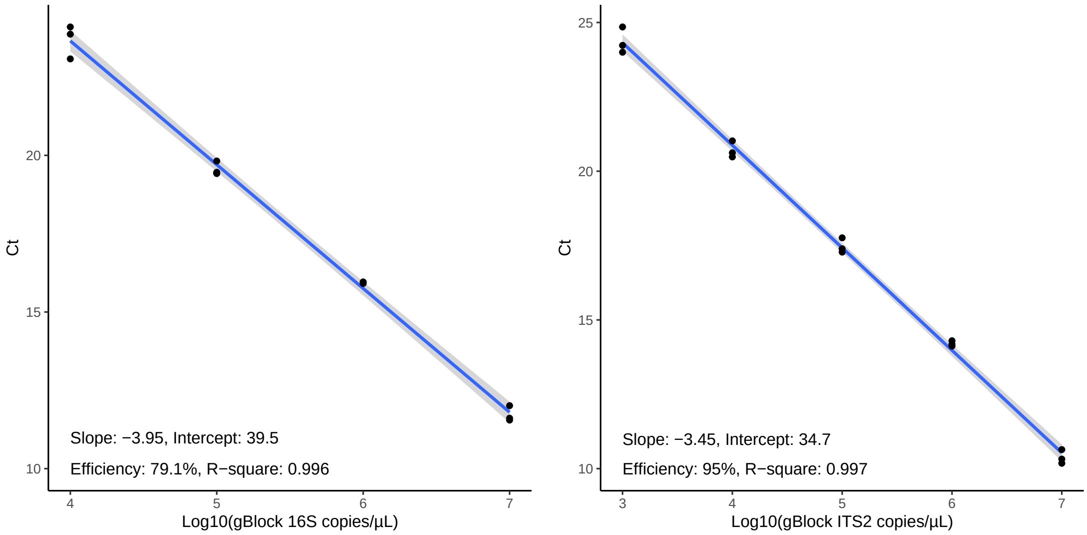


(a) Bacteria

(b) Fungi

Figure S1: **Standard curve for qPCR analysis of bacteria (16S V3-V4) and fungi (ITS2).** A synthetic double-stranded DNA fragment (G-block) containing a single copy of the target gene sequence was used to prepare the standard dilution series ranging from $10^7$ to $10^1$ copies. The standards $10^1$ to $10^3$ for bacteria and $10^1$ to $10^2$ for fungi were excluded from the analysis due to the presence of primer dimer signals, which became apparent at these higher dilutions. For bacterial, all samples amplified within the standard range of $10^5$ to $10^7$, while fungal samples within the range of $10^3$ to $10^5$.

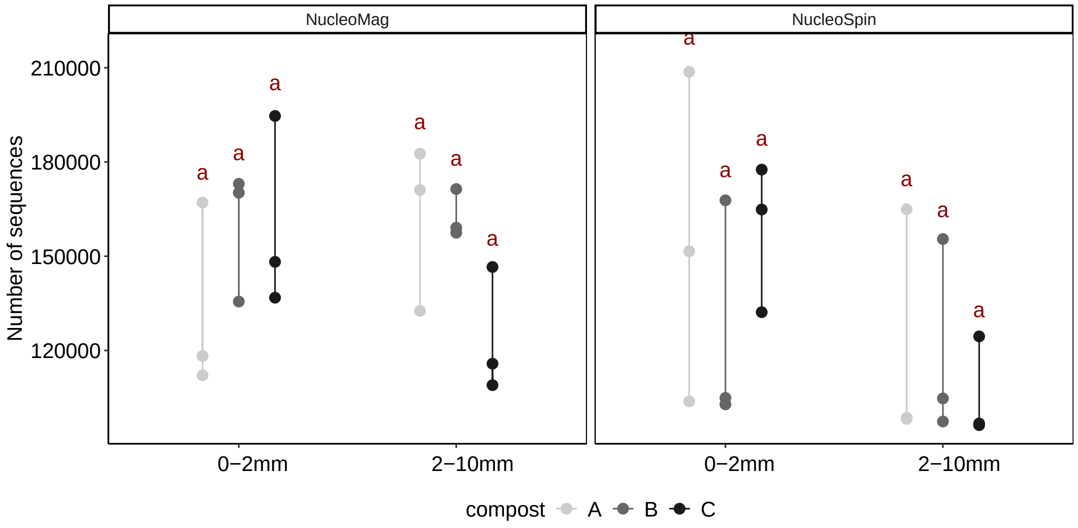


(a) Bacteria

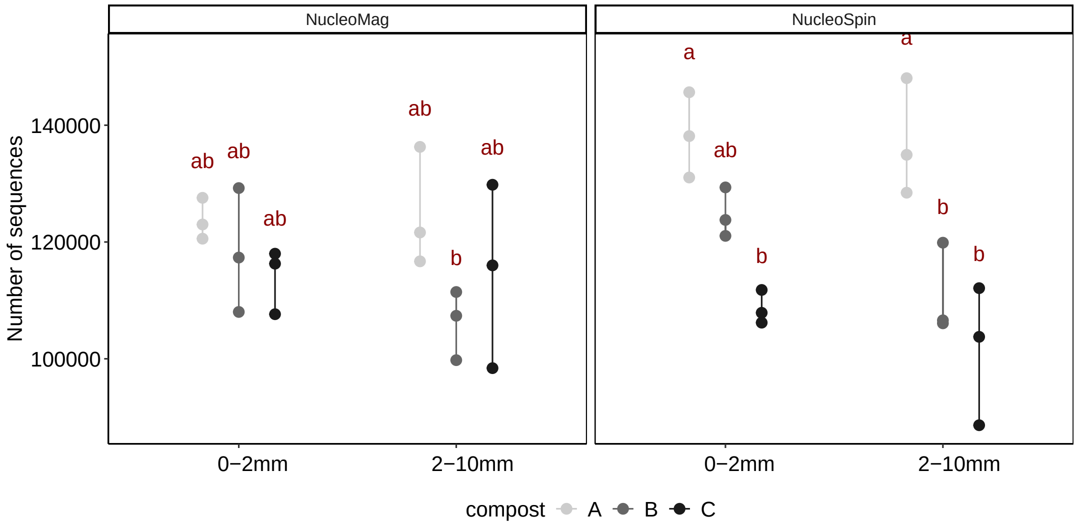


(b) Fungi

Figure S2: **The number of sequences varied between treatments for fungi, but not for bacteria.** (b) Fungal sequences varied across composts, with a stronger differences when extracted with the column-based than with the bead-based method (ANOVA results: $F_{Compost} = 19.1$, $p$ <0.001; $F_{Compost:Kit} = 5.2$, $p = 0.014$)

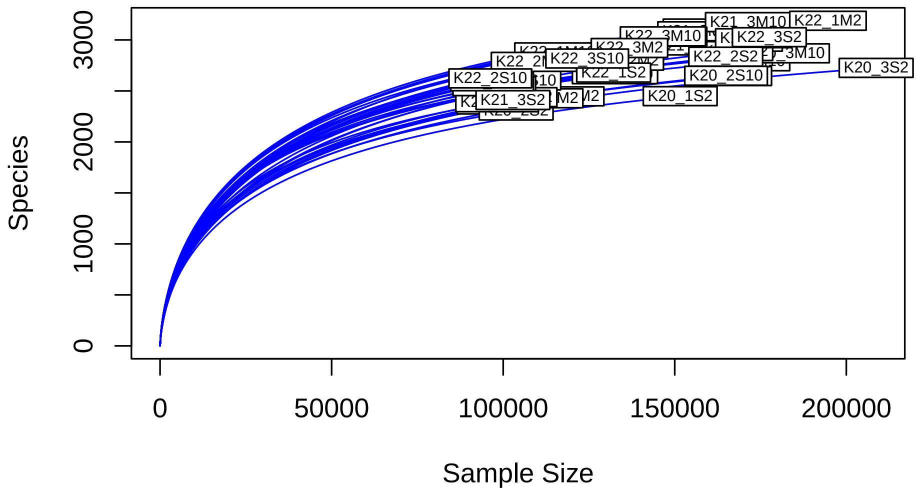


(a) Bacteria

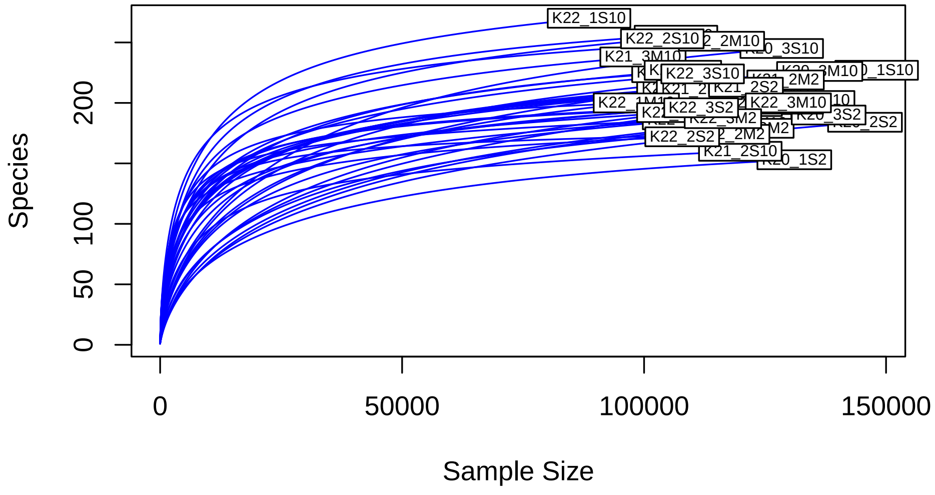


(b) Fungi

Figure S3: The flattening of rarefaction curves shows that both (a) bacterial and (b) fungal diversity is well covered in the samples.

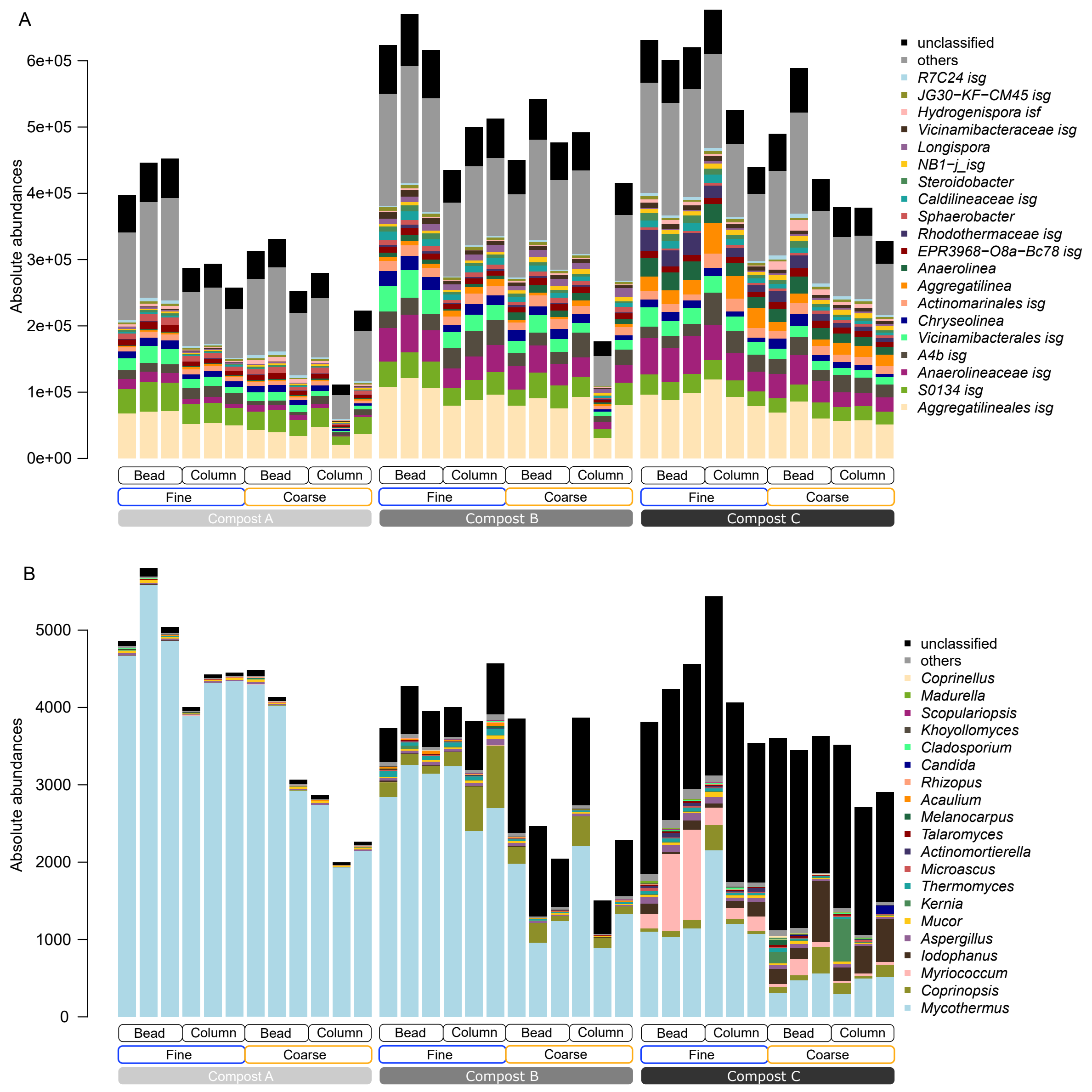


Figure S4: **Similar most abundant taxa are found in compost A-C, but differ in their abundance.** Barplots show absolute abundances of (A) bacteria and (B) fungi for all 36 samples. The three technical replicates are shown next to each other. Absolute abundances were calculated by multiplying the relative abundances (based on unrarefied reads) with the target gene copy number assessed by qPCR. The abbreviations "isg" and "isf" stand for *incertae sedis* genus and family, respectively, which means that the systematic of the taxa is unknown at higher resolution.

Table S1: Primer sequences used for quantitative real-time PCR and amplicon sequencing for fungi and bacteria. Degenerated bases are shown in bold. Amplicon lenght for bacteria 4̃60 bp, and for fungi 250-255 bp.

| Analysis | name | sequence | reference |
|---|---|---|---|
| Bacteria qPCR | 341F | 5’-CCTACGGG**D**GGC**W**GCA-3’ | Liu et al. (2012) |
| | 806R | 5’-GGACTAC**HV**GGGT**M**TCTAATC-3’ | |
| Bacteria sequencing | 341F | 5’-CCTA**Y**GGG**DB**GC**WS**CAG-3’ | Frey et al. (2016) |
| | 806R | 5’-GGACTAC**NV**GGGT**H**TCTAAT-3’ | |
| Fungi qPCR | ITS3 | 5’-CA**H**CGATGAAGAACG**YR**G-3’ | Frey et al. (2016) |
| | ITS4 | 5’-TCCT**S**CGCTTATTGATATGC-3’ | |
| Fungi sequencing | ITS3 | 5’-CA**N**CGATGAAGAACG**YR**G-3’ | Tedersoo et al. (2015) |
| | ITS4 | 5’-CCT**S**C**S**CTTA**N**T**D**ATATGC-3’ | |

Table S2: The table displays for all 36 samples the DNA concentration of DNA extracts, $\log_{10}$ of 16S and ITS2 copy numbers normalized by the qPCR standard curve and the absolute copy numbers normliazed by DNA concentration.

| comp | size | kit | rep | DNA conc [ng/qL] | log(16S copies) | 16S copies [1/ng DNA] | log(ITS2 copies) | ITS copies [1/ng DNA] |
|---|---|---|---|---|---|---|---|---|
| A | 2 | M | 1 | 24.8 | 5.99 | 397122 | 4.08 | 4859 |
| A | 2 | M | 2 | 28.3 | 6.10 | 446182 | 4.22 | 5800 |
| A | 2 | M | 3 | 26.9 | 6.09 | 452086 | 4.13 | 5035 |
| A | 2 | S | 1 | 18.0 | 5.71 | 287113 | 3.86 | 4003 |
| A | 2 | S | 2 | 24.5 | 5.86 | 293296 | 4.04 | 4426 |
| A | 2 | S | 3 | 19.8 | 5.71 | 257854 | 3.94 | 4452 |
| A | 10 | M | 1 | 9.8 | 5.49 | 313045 | 3.64 | 4481 |
| A | 10 | M | 2 | 12.6 | 5.62 | 331342 | 3.72 | 4134 |
| A | 10 | M | 3 | 12.8 | 5.51 | 252973 | 3.59 | 3066 |
| A | 10 | S | 1 | 12.1 | 5.53 | 279442 | 3.54 | 2866 |
| A | 10 | S | 2 | 10.6 | 5.07 | 111485 | 3.33 | 1996 |
| A | 10 | S | 3 | 11.8 | 5.42 | 223076 | 3.42 | 2264 |
| B | 2 | M | 1 | 47.3 | 6.47 | 623304 | 4.25 | 3731 |
| B | 2 | M | 2 | 36.9 | 6.39 | 669933 | 4.20 | 4281 |
| B | 2 | M | 3 | 36.5 | 6.35 | 615511 | 4.16 | 3949 |
| B | 2 | S | 1 | 21.1 | 5.96 | 434867 | 3.93 | 4005 |
| B | 2 | S | 2 | 34.3 | 6.23 | 499888 | 4.12 | 3817 |
| B | 2 | S | 3 | 24.0 | 6.09 | 512789 | 4.04 | 4569 |
| B | 10 | M | 1 | 30.4 | 6.14 | 449904 | 4.07 | 3858 |
| B | 10 | M | 2 | 41.8 | 6.36 | 542439 | 4.01 | 2467 |
| B | 10 | M | 3 | 47.5 | 6.36 | 476606 | 3.99 | 2046 |
| B | 10 | S | 1 | 25.5 | 6.10 | 491861 | 3.99 | 3871 |
| B | 10 | S | 2 | 25.1 | 5.65 | 176259 | 3.58 | 1502 |
| B | 10 | S | 3 | 25.3 | 6.02 | 415581 | 3.76 | 2281 |
| C | 2 | M | 1 | 23.9 | 6.18 | 631543 | 3.96 | 3815 |
| C | 2 | M | 2 | 28.4 | 6.23 | 600516 | 4.08 | 4238 |
| C | 2 | M | 3 | 22.0 | 6.13 | 620321 | 4.00 | 4561 |
| C | 2 | S | 1 | 34.3 | 6.37 | 676754 | 4.27 | 5439 |
| C | 2 | S | 2 | 24.2 | 6.10 | 524802 | 3.99 | 4064 |
| C | 2 | S | 3 | 21.4 | 5.97 | 439162 | 3.88 | 3543 |
| C | 10 | M | 1 | 28.1 | 6.14 | 489930 | 4.00 | 3598 |
| C | 10 | M | 2 | 41.0 | 6.38 | 588453 | 4.15 | 3444 |
| C | 10 | M | 3 | 36.6 | 6.19 | 421025 | 4.12 | 3633 |
| C | 10 | S | 1 | 17.3 | 5.82 | 378798 | 3.78 | 3517 |
| C | 10 | S | 2 | 18.0 | 5.83 | 377949 | 3.69 | 2707 |
| C | 10 | S | 3 | 18.8 | 5.79 | 328326 | 3.74 | 2905 |

Table S3: Number of sequences and ASVs for the different filtering steps of sequencing data.

| **Group** | **Step** | **Type** | **Total** | **Min** | **Max** | **Mean** |
|---|---|---|---|---|---|---|
| Bacteria | High-quality | Sequences | 5116329 | 97659 | 211035 | 142120 |
| | Only bacteria | | 5052447 | 96235 | 208697 | 140346 |
| | Rarefied | | 3474088 | 96102 | 96744 | 96502 |
| | High-quality | ASVs | 4947 | 2334 | 3242 | 2792 |
| | Only bacteria | | 4871 | 2308 | 3188 | 2754 |
| | Rarefied | | 4871 | 2308 | 3114 | 2720 |
| Fungi | High-quality | Sequences | 4484713 | 99323 | 154607 | 124575 |
| | Only fungi | | 4268339 | 88625 | 148053 | 118565 |
| | Rarefied | | 3191333 | 88625 | 88674 | 88648 |
| | High-quality | ASVs | 1752 | 417 | 743 | 545 |
| | Only fungi | | 851 | 153 | 270 | 207 |
| | Rarefied | | 851 | 153 | 270 | 207 |

Table S4: Number of treatment-specific ASVs for particle size fraction and DNA extraction method for each compost separately and shared between composts. In brackets the summed mean relative abundance across all samples of the respective compost is indicated. The analysis is based on robustly detected ASVs (at least present in 2 out of 3 biological replicates for one treatment).

| | **compost** | **A** | **B** | **C** | **AB** | **BC** | **AC** | **ABC** |
|---|---|---|---|---|---|---|---|---|
| Bacteria Particle size | **both** | 3326 (99.86) | 3810 (99.87) | 3888 (99.85) | | | | |
| | **fine** | 238 (0.04) | 203 (0.04) | 234 (0.07) | 11 | 21 | 12 | 1 |
| | **coarse** | 323 (0.09) | 314 (0.09) | 253 (0.09) | 36 | 42 | 24 | 8 |
| Bacteria DNA extraction | **both** | 3347 (99.89) | 3838 (99.89) | 3923 (99.88) | | | | |
| | **column** | 217 (0.04) | 177 (0.04) | 190 (0.05) | 15 | 21 | 12 | 5 |
| | **beads** | 323 (0.07) | 312 (0.07) | 262 (0.07) | 33 | 44 | 30 | 5 |
| Fungi Particle size | **both** | 315 (99.97) | 306 (99.87) | 286 (99.78) | | | | |
| | **fine** | 26 (0.01) | 47 (0.07) | 32 (0.06) | 3 | 8 | 7 | 1 |
| | **coarse** | 51 (0.03) | 49 (0.06) | 77 (0.15) | 8 | 13 | 16 | 4 |
| Fungi DNA extraction | **both** | 316 (99.97) | 309 (99.86) | 291 (99.77) | | | | |
| | **column** | 38 (0.02) | 59(0.09) | 59 (0.13) | 12 | 16 | 10 | 4 |
| | **beads** | 38 (0.01) | 34(0.04) | 45 (0.10) | 6 | 3 | 9 | 0 |

Table S5: Number of treatment-enriched ASVs for particle size fraction and DNA extraction method for each compost separately and shared between compost. In brackets the summed mean relative abundance across all samples of the respective compost is indicated. The analysis is based on absolute abundances (relative abundance of unrarefied reads multiplied with target gene copy number).

| | compost | A | B | C | AB | BC | AC | ABC |
|---|---|---|---|---|---|---|---|---|
| Bacteria | **fine** | 62 (2.99) | 48 (5.50) | 93 (8.60) | 3 | 24 | 8 | 1 |
| Particle size | **coarse** | 28 (1.20) | 21 (0.38) | 47 (2.70) | 1 | 8 | 4 | 1 |
| Bacteria | **column** | 1 (0.01) | 0 | 5 (0.05) | 0 | 0 | 0 | 0 |
| DNA extraction | **beads** | 103 (4.94) | 42 (1.64) | 61 (2.09) | 7 | 10 | 5 | 1 |
| Fungi | **fine** | 0 | 2 (64.46) | 5 (41.67) | 0 | 1 | 0 | 0 |
| Particle size | **coarse** | 0 | 1 (12.42) | 1 (6.95) | 0 | 0 | 0 | 0 |
| Fungi | **column** | 0 | 0 | 0 | 0 | 0 | 0 | 0 |
| DNA extraction | **beads** | 0 | 0 | 0 | 0 | 0 | 0 | 0 |

Table S6: Analysis of Variance Table for the effect of compost, particle size fraction (size), and DNA extraction method (kit) on bacterial quantity and bacterial and fungal evenness. For this variables the effect of the "compost_replicate" was close to zero and therefore a standard linear model were used.

| | | df | Sums of Sq | Mean of Sq | F-value | p-value |
|---|---|---|---|---|---|---|
| **Bacterial quantity** | compost | 1 | 3.08E+11 | 1.54E+11 | 28.43 | 0.000 |
| | kit | 2 | 1.36E+11 | 1.36E+11 | 25.15 | 0.000 |
| | size | 2 | 1.51E+11 | 1.51E+11 | 28.00 | 0.000 |
| | compost:kit | 3 | 2.03E+09 | 1.01E+09 | 0.19 | 0.830 |
| | compost:size | 2 | 3.50E+09 | 1.75E+09 | 0.32 | 0.727 |
| | kit:size | 3 | 6.15E+07 | 6.15E+07 | 0.01 | 0.916 |
| | compost:kit:size | 2 | 6.36E+09 | 3.18E+09 | 0.59 | 0.563 |
| | Residuals | 24 | 1.30E+11 | 5.41E+09 | NA | NA |
| **Bacterial evenness** | compost | 2 | 1.40E-03 | 7.00E-04 | 21.12 | 0.000 |
| | kit | 1 | 2.35E-03 | 2.35E-03 | 70.80 | 0.000 |
| | size | 1 | 4.24E-03 | 4.24E-03 | 127.91 | 0.000 |
| | compost:kit | 2 | 5.27E-04 | 2.63E-04 | 7.94 | 0.002 |
| | compost:size | 2 | 2.35E-04 | 1.18E-04 | 3.55 | 0.045 |
| | kit:size | 1 | 3.90E-06 | 3.90E-06 | 0.12 | 0.735 |
| | compost:kit:size | 2 | 2.06E-04 | 1.03E-04 | 3.10 | 0.063 |
| | Residuals | 24 | 7.96E-04 | 3.32E-05 | NA | NA |
| **Fungal evenness** | compost | 2 | 8.59E-01 | 4.30E-01 | 542.39 | 0.000 |
| | kit | 1 | 1.01E-03 | 1.01E-03 | 1.28 | 0.270 |
| | size | 1 | 1.38E-03 | 1.38E-03 | 1.75 | 0.199 |
| | compost:kit | 2 | 4.38E-03 | 2.19E-03 | 2.77 | 0.083 |
| | compost:size | 2 | 1.19E-02 | 5.95E-03 | 7.52 | 0.003 |
| | kit:size | 1 | 1.58E-04 | 1.58E-04 | 0.20 | 0.660 |
| | compost:kit:size | 2 | 5.75E-03 | 2.87E-03 | 3.63 | 0.042 |
| | Residuals | 24 | 1.90E-02 | 7.92E-04 | NA | NA |

Table S7: Analysis of Deviance Table (Type III Wald chisquare tests) for the effect of compost, particle size fraction (size), and DNA extraction method (kit) on bacterial richness and fungal quantity and fungal richness. Mixed linear models with "compost_replicate" as a random factor was used.

| | | **Chisq** | **Df** | **Pr(<Chisq)** |
|---|---|---|---|---|
| **Bacterial richness** | Intercept | 12508.99 | 1 | 0.000 |
| | size | 174.14 | 2 | 0.000 |
| | compost | 10.66 | 1 | 0.001 |
| | kit | 32.58 | 1 | 0.000 |
| | compost:kit | 9.27 | 2 | 0.010 |
| | compost:size | 18.40 | 2 | 0.000 |
| | kit:size | 1.68 | 1 | 0.195 |
| | compost:kit:size | 5.84 | 2 | 0.054 |
| **Fungal quantity** | Intercept | 299.66 | 1 | 0.000 |
| | size | 9.67 | 2 | 0.008 |
| | compost | 4.84 | 1 | 0.028 |
| | kit | 9.85 | 1 | 0.002 |
| | compost:kit | 4.29 | 2 | 0.117 |
| | compost:size | 1.47 | 2 | 0.480 |
| | kit:size | 0.93 | 1 | 0.335 |
| | compost:kit:size | 0.11 | 2 | 0.945 |
| **Fungal richness** | Intercept | 289.02 | 1 | 0.000 |
| | size | 5.42 | 2 | 0.067 |
| | compost | 2.71 | 1 | 0.100 |
| | kit | 0.09 | 1 | 0.764 |
| | compost:kit | 2.11 | 2 | 0.349 |
| | compost:size | 2.67 | 2 | 0.263 |
| | kit:size | 4.57 | 1 | 0.032 |
| | compost:kit:size | 1.55 | 2 | 0.461 |

Table S8: PERMANOVA results testing the effect of compost, particle size fraction (size) and DNA extraction method (kit) on (a) bacterial and (b) fungal community structure (based on Bray-Curtis dissimilarities).

(a) Bacteria

| Factor | Df | $R^2$ [%] | Pseudo-F | $p$ |
|---|---|---|---|---|
| compost | 2 | 75.9 | 142.7 | 0.001 |
| size | 1 | 6.3 | 23.8 | 0.001 |
| kit | 1 | 6.0 | 22.4 | 0.001 |
| compost:size | 2 | 3.0 | 5.7 | 0.002 |
| compost:kit | 2 | 0.8 | 1.5 | 0.23 |
| size:kit | 1 | 0.8 | 3.0 | 0.055 |
| compost:size:kit | 2 | 0.8 | 1.6 | 0.161 |
| residual | 24 | 6.4 | | |

(b) Fungi

| Factor | Df | $R^2$ [%] | Pseudo-F | $p$ |
|---|---|---|---|---|
| compost | 2 | 71.0 | 105.3 | 0.001 |
| size | 1 | 8.9 | 26.4 | 0.001 |
| kit | 1 | 0.8 | 2.2 | 0.115 |
| compost:size | 2 | 8.8 | 13.1 | 0.001 |
| compost:kit | 2 | 1.3 | 1.9 | 0.111 |
| size:kit | 1 | 0.4 | 1.3 | 0.263 |
| compost:size:kit | 2 | 0.7 | 1.0 | 0.382 |
| residual | 24 | 8.1 | | |
